# N-acetylcysteine counteracts immune dysfunction and autistic-related behaviors in the *Shank3b* mouse model of Autism Spectrum Disorders

**DOI:** 10.1101/2024.03.13.584809

**Authors:** Luca Pangrazzi, Enrica Cerilli, Luigi Balasco, Ginevra Matilde Dall’O’, Gabriele Chelini, Anna Pastore, Birgit Weinberger, Yuri Bozzi

## Abstract

Autism Spectrum Disorder (ASD) includes a range of neurodevelopmental disabilities characterized by social interaction deficits, communication impairments, and repetitive behaviors. Previous studies have shown that pro-inflammatory conditions play a key role in ASD. Here we reported that increased levels of molecules related to inflammation are present in the cerebellum and peripheral blood (PB) of mice lacking Shank3b, established model of syndromic ASD. In parallel, immune dysfunction was documented in the bone marrow (BM) and spleen of mutant mice. N-acetylcysteine (NAC) treatment rescued inflammation in the cerebellum and PB, as well as impaired production of pro-inflammatory molecules in the BM and spleen. In addition, social impairment was counteracted in NAC-treated Shank3b^-/-^ animals. Taken together, our study further confirms the key role of cerebellar inflammation in the establishment of ASD-related behaviors. Furthermore, our findings underscore the importance of considering ASD as a systemic disorder.

Our findings therefore suggest that the interplay between oxidative stress and inflammation may support ASD-related behaviors in mice.

## Introduction

Autism Spectrum Disorder (ASD) identifies a group of neurodevelopmental disabilities associated with impairments in social interaction as well as social communication, repetitive behaviors and interests (1). Alongside with the core symptoms, comorbidities including motor impairments, epilepsy, anxiety, depression, obsessive-compulsive disorder (OCD), and attention-deficit/hyperactivity disorder (ADHD) can be found in ASD (2). The estimated worldwide prevalence of ASD is around 1/100 children, although it may be even higher in some countries (3,4). ASD pathophysiology is very complex, as both genetic and environmental factors play an important role (5). In the last 20 years, a consistent amount of work has been done to identify genes associated with ASD and strategies of intervention have been proposed (6). Despite this, effective medical treatments have not been identified yet.

Immune dysfunction and inflammation have been proposed as key contributors to the pathogenesis and the severity of ASD (7). In humans, increased levels of pro-inflammatory molecules tumour necrosis factor (TNF), interleukin (IL)-1β, IL-6, IL-8, IL-12p40, and IL-17 were described in the plasma of ASD children as compared with age-matched healthy controls (8,9). Furthermore, cytokines IL-1β, IL-6, TNF, interferon (IFN)γ, and chemokine C-C motif ligand (CCL)-2) were overexpressed in the cerebrospinal fluid and brain of ASD subjects (10,11). Similarly, increased levels of pro-inflammatory molecules in brain areas and in the peripheral blood (PB) of ASD mouse models have been reported (7). In particular, our recent work described pro-inflammatory dysfunction in the cerebellum and in the periphery of mice lacking the *Cntnap2* gene (*Cntnap2^-/-^*mice; 12), a well-established model of syndromic ASD.

Accumulation of oxygen radicals, a condition commonly known as oxidative stress, can generally be found in the presence of inflammation. Increased ROS levels, paralleled by decreased antioxidant capacity, have been described in both ASD and mouse models (13–16). Transcriptomic analyses performed in mice showed that genes coding for ROS scavenging enzymes were less expressed in the brain of mouse models of ASD (7). In *Cntnap2^-/-^* mice, we showed that systemic pro-inflammatory processes and ASD-related behaviors could be counteracted by the chronic administration of the antioxidant/anti-inflammatory molecule N-acetylcysteine (NAC; 12). Thus, strong evidence exists that a connection between oxidative stress and inflammation may play a determinant role in ASD.

Mice lacking the SH3 and multiple ankyrin repeat domains protein 3b (*Shank3b*^-/-^ mice) represent a well-established mouse model of ASD, as they show deficits in social interaction, self-injurious repetitive grooming, and impaired locomotor activity (17, 18). In humans, SHANK3 mutations are responsible for the development of 22q13 deletion syndrome (Phelan-McDermid Syndrome) and other non-syndromic forms of ASD (19).

In this study, we measured the expression of molecules related to inflammation in the cerebral cortex, hippocampus, cerebellum, and PB of *Shank3b*^-/-^, *Shank3b*^+/-^, and *Shank3b*^+/+^ mice. In parallel, immune parameters were assessed in the bone marrow (BM) and spleen, lymphoid organs respectively involved in the production/maintenance of immune cells and in the generation of immune responses. Pro-inflammatory dysfunction was detected in the cerebellum and PB of mutant animals. Furthermore, a differential expression of molecules involved into pro-inflammatory processes was additionally documented in the BM and spleen. Finally, NAC treatments could rescue social deficits, repetitive behaviors, and immune dysfunction in mutant mice. Altogether this work strengthens the evidence that pro-inflammatory processes within the cerebellum may support ASD-related behaviors in mice.

## Results

### Molecules related to inflammation increase in the cerebellum of mutant ***Shank3b* mice**

To assess whether inflammation may be present in the *Shank3b* model, the expression of key pro-inflammatory molecules was measured in the cerebral cortex, cerebellum and hippocampus of *Shank3b*^+/+^, *Shank3b*^+/-^, and *Shank3b*^-/-^ mice (Fig. 1). Overall, the levels of all molecules related to inflammation were increased in the cerebellum in comparison with the other brain areas (Fig. 1 a-g). TNF expression within the cerebellum was the highest in *Shank3b*^-/-^ animals, while no differences were observed between *Shank3b*^+/-^ and *Shank3b*^+/+^ mice (Fig. 1a). These results were confirmed using flow cytometry, which described increased TNF levels in CD14^+^ cerebellar cells from *Shank3b*^-/-^ animals. Gating strategy used for the flow cytometry experiments is shown in Suppl. Fig. 1. IFNγ expression was detected in the cerebellum, but not in the cerebral cortex and in the hippocampus (Fig. 1b). Again, IFNγ levels were the highest in *Shank3b*^-/-^ and the lowest in *Shank3b*^+/+^ mice, while in this case they were intermediate in the *Shank3b*^+/-^ group. The same results were described at the protein level when IFNγ expression was measured within T cells. Similar trends were observed for the pro-inflammatory molecules IL-6 and IL-1β, as their mRNAs were overexpressed in the cerebellum of *Shank3b*^-/-^ animals (Fig. 1 c-d). Chemokines CCL3, CCL5, and CCL20 control the chemotaxis of immune cells to sites of inflammation (20). Again, mRNAs of all three molecules could be measured in the cerebellum only (Fig. 1 e-g). Also in this case, the expression of these chemokines was high in the *Shank3b*^-/-^ group, intermediate in *Shank3b*^+/-^ mice (at least for CCL3 and CCL20 mRNAs) and low in *Shank3b*^+/+^ controls. mRNA levels of metalloprotease (MMP) 8, known to support tissue damage and to play a determinant role in neuroinflammation (21), were increased within the cerebellum and again were the highest in *Shank3b*^-/-^ mice (Fig. 1h). Inhibitor of cyclin-dependent kinases p21 (CDKN1A), activated during DNA damage response and additionally involved in cellular senescence (22), showed a similar mRNA expression in the cerebral cortex and in the cerebellum, while it was reduced in the hippocampus (Fig. 1i). Interestingly, p21 mRNA levels within the cerebellum were low in the *Shank3b*^+/+^ group, intermediate in *Shank3b*^+/-^ and high in mutant animals. We next investigated whether pro-inflammatory changes may additionally be present in the peripheral blood (PB). Indeed, the expression of TNF in CD14^+^ cells and IFNγ within CD8^+^ and CD4^+^ T cells was increased in both *Shank3b*^+/-^ and *Shank3b*^-/-^ animals, in comparison with the control group (Figure 1 j-l). Taken together, pro-inflammatory impairments are present in the cerebellum and in the PB of *Shank3b*^-/-^ and *Shank3b*^+/-^ mice.

**Figure 1.**
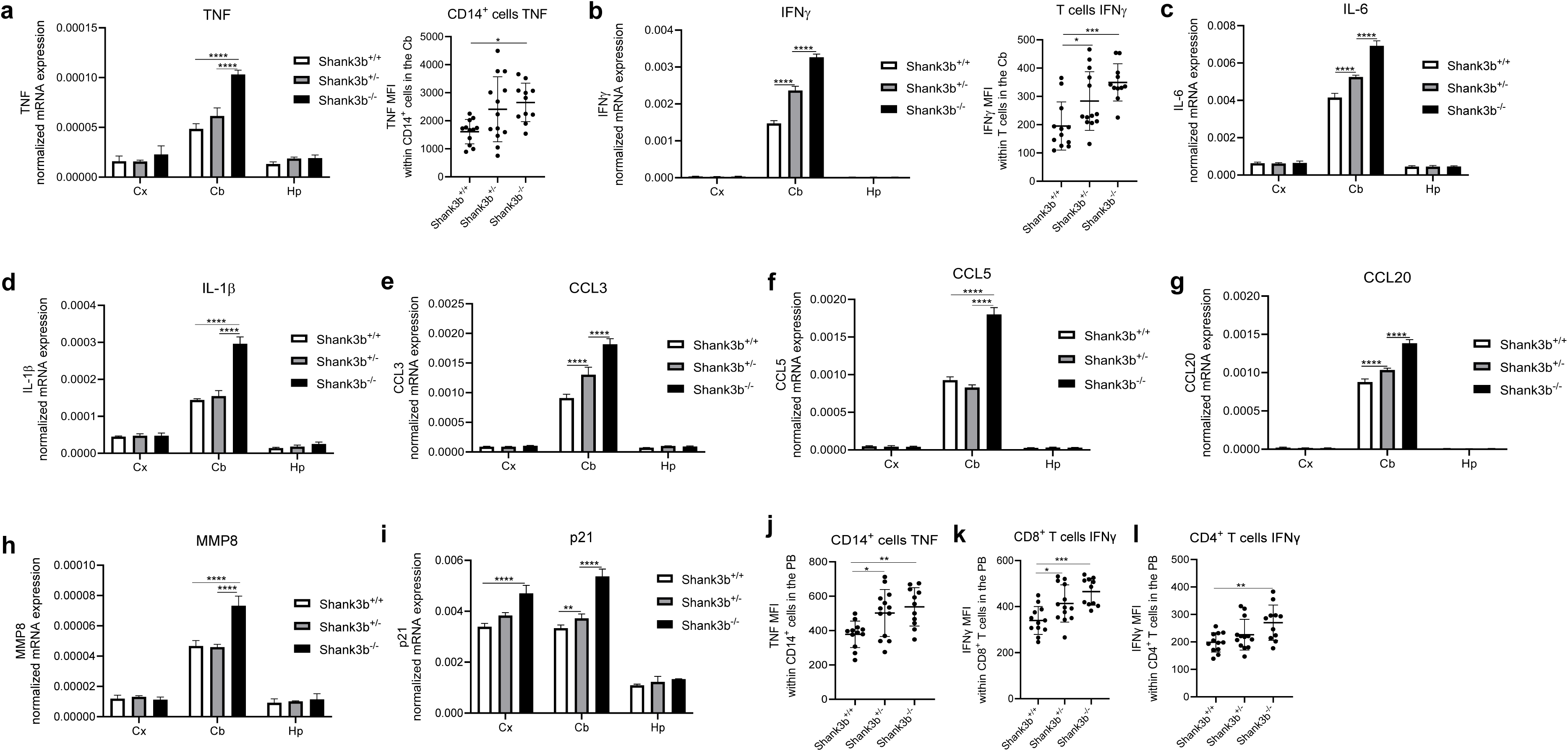
Pro-inflammatory changes in the brain of *Shank3b^+/+^*, *Shank3b^+/-^* and *Shank3b^-/-^* mice. (**a**) TNF mRNA expression in the cerebral cortex (Cx), cerebellum (Cb) and hippocampus (Hp) of *Shank3b^+/+^, Shank3b^+/-^* and *Shank3b^-/-^*mice measured using RT-qPCR, and TNF expression at the protein level (shown as mean fluorescence intensity, MFI) within cerebellar CD14^+^ cells assessed using flow cytometry. (**b**) IFNγ mRNA expression in the Cx, Cb, and Hp and IFNγ levels within cerebellar T cells. mRNA expression of (**c**) IL-6, (**d**) IL-1β, I CCL3, (**f**) CCL5, (**g**) CCL20, (**h**) MMP8, (**i**) p21 in the Cx, Cb, and Hp. mRNA expression was normalized against housekeeping gene β-actin. (**j**) TNF MFI within CD14^+^ cells, IFNγ MFI within (**k**) CD8^+^ T cells and (**l**) CD4^+^ T cells in the PB of *Shank3b^+/+^, Shank3b^+/-^* and *Shank3b^-/-^* mice. One-way ANOVA, Tukey post-hoc test. N = 8 per group (RT-qPCR); n = 10-12 per group (flow cytometry). *p<0.05; **p<0.01; ***p<0.001; ****p<0.0001.

### Immune dysfunction is present in the bone marrow and spleen of mutant ***Shank3b* mice**

As a next step, we assessed whether immune system dysfunction may additionally be observed in the bone marrow (BM) and spleen of *Shank3b*^+/-^ and *Shank3b*^-/-^ mice (Fig. 2). In both organs, the expression of pro-inflammatory molecules varied consistently across genotypes. Within the BM, although TNF was overexpressed in *Shank3b*^-/-^ animals in comparison to the control group, the levels of this molecule were reduced in *Shank3b*^+/-^ mice (Fig. 2a). No differences were described in the spleen. These results were confirmed at the protein level using flow cytometry, in which TNF expression was assessed in CD14^+^ cells, a population including macrophages and monocytes. No differences were found within CD8^+^ and CD4^+^ T cells, in both BM and spleen (Fig. 2a and Suppl. Fig.2 a-d). Surprisingly, IFNγ mRNA levels in the BM were reduced in both *Shank3b*^+/-^ and *Shank3b*^-/-^ mice, and in the spleen of *Shank3b*^-/-^ animals, when compared with the *Shank3b*^+/+^ group (Fig. 2b). Similar results were obtained when IFNγ expression was measured within CD8^+^ T cells and in CD4^+^ T cells, although no differences between *Shank3b*^+/+^ and *Shank3b*^+/-^ were observed (Fig. 2b and Suppl. Fig. 2 e,f). Furthermore, comparable IL-6 levels were detected in *Shank3b*^+/+^ and *Shank3b*^-/-^ mice in the BM, while the expression of this cytokine was again reduced in *Shank3b*^+/-^ mice (Fig. 2c). Despite this, increased IL-6 levels in *Shank3b*^-/-^ mice and similar expression between the other two groups were observed within BM CD14^+^ cells. Very low IL-6 levels were found within T cells (data not shown). In addition, no differences were identified in the spleen. IL-1β mRNA expression was again increased in the BM of *Shank3b*^-/-^ and decreased in *Shank3b*^+/-^ mice in comparison with the control group, while no differences were found in the spleen (Fig. 2d). We next investigated the mRNA expression of chemokines CCL3, CCL5 and CCL20 in the BM and in the spleen (Fig. e-g). Interestingly, when *Shank3b*^+/+^ and *Shank3b*^-/-^ animals were compared, no differences for CCL3 and CCL8 and decreased CCL20 mRNA levels were described in the BM of mutant mice. In the spleen, reduced CCL5 and CCL20 mRNA levels were found in *Shank3b*^-/-^ animals when compared with controls. In parallel, decreased CCL3 and CCL20 mRNAs in the BM, reduced CCL5 mRNA and increased CCL20 mRNA in the spleen were described in *Shank3b*^+/-^ animals compared with the *Shank3b*^+/+^ group. Thus, these results suggest that the migratory capacity of immune cells in the BM and in the spleen may be different between *Shank3b*^+/+^, *Shank3b*^+/-^, and *Shank3b*^-/-^ mice. Similarly, MMP8 mRNA levels were decreased in the BM of *Shank3b*^+/-^ and *Shank3b*^-/-^ animals, reduced in the spleen of *Shank3b*^-/-^ and increased in the *Shank3b*^+/-^ group (Fig. 2i). Furthermore, p21 mRNA expression in the BM was upregulated in *Shank3b*^-/-^ and downregulated in *Shank3b*^-/+^ mice when compared with control animals, while no differences were present in the spleen (Fig. 2i). IL-15 is known to play a determinant role in the maintenance of immune cells in the BM and in the establishment of immunological memory in the spleen (23,24). mRNA levels of this cytokine in *Shank3b*^-/-^ mice was found to be reduced in the BM and increased in the spleen as compared with the other two groups. Finally, we observed that mRNA expression of the antioxidant enzyme SOD3 was decreased in the BM and spleen of *Shank3b*^-/-^ animals and, in the BM, also in *Shank3b*^+/-^ mice (Fig. 2k). Taken together, the levels of molecules related to inflammation in the BM and spleen change in *Shank3b*^+/-^ and *Shank3b*^-/-^ mice. In *Shank3b*^-/-^ mice, while pro-inflammatory processes are induced in the BM, they may be inhibited in the spleen. Furthermore, these impairments may be accompanied by oxidative stress.

**Figure 2.**
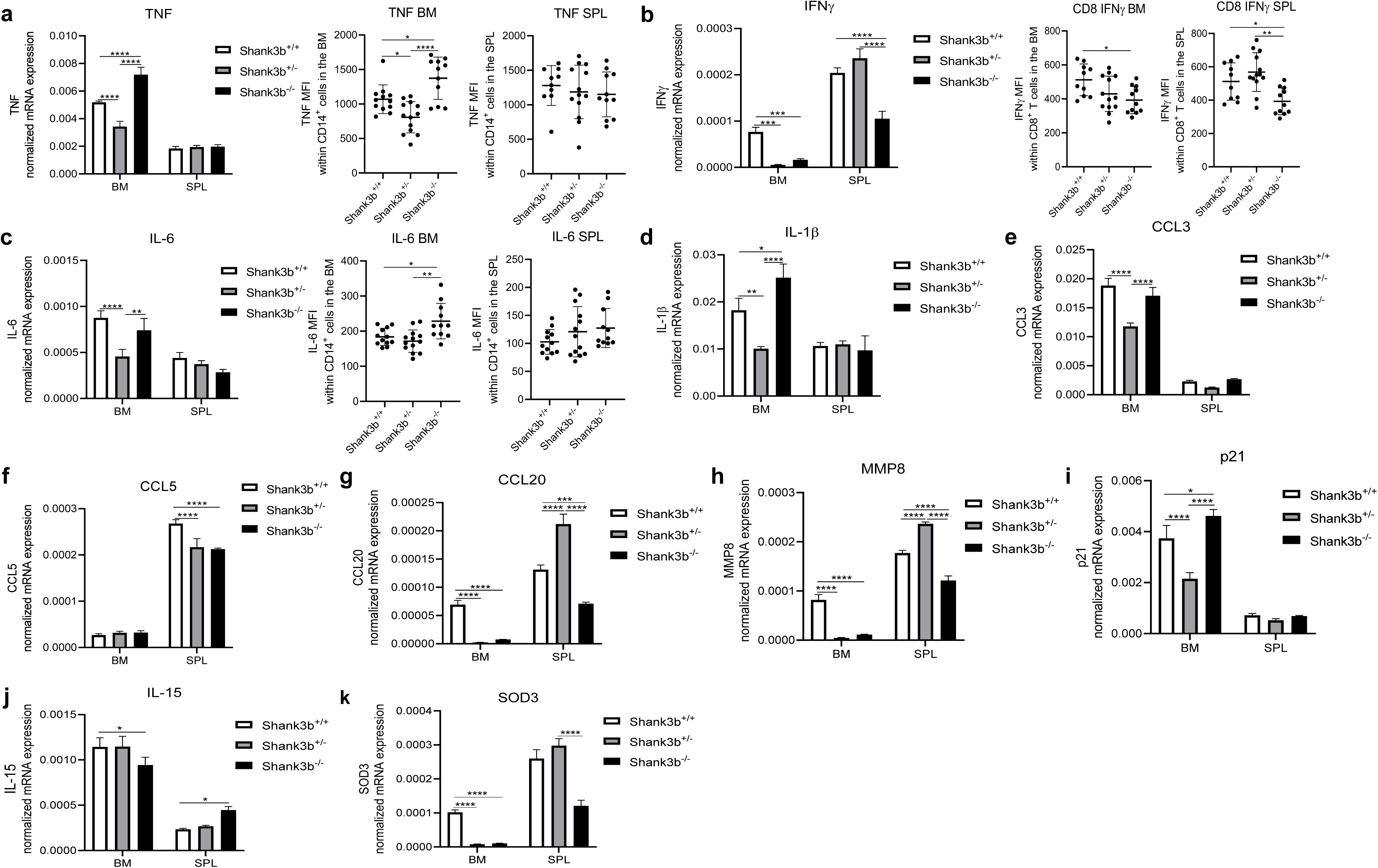
Pro-inflammatory changes in the bone marrow and spleen of *Shank3b^+/+^*, *Shank3b^+/-^* and *Shank3b^-/-^* mice. (**a**) TNF mRNA expression in the bone marrow (BM) and spleen (SPL) of *Shank3b^+/+^, Shank3b^+/-^* and *Shank3b^-/-^* mice measured using RT-qPCR, and TNF expression at the protein level (shown as mean fluorescence intensity, MFI) within CD14^+^ cells assessed using flow cytometry. (**b**) IFNγ mRNA expression and IFNγ levels within CD8^+^ T cells. (**c**) IL-6 mRNA expression and IL-6 levels within CD14^+^ cells. mRNA expression of (**d**) IL-1β, (**e**) CCL3, (**f**) CCL5, (**g**) CCL20, (**h**) MMP8, (**i**) p21, (**j**) IL-15, and (**k**) SOD3 in the BM and SPL. mRNA expression was normalized against housekeeping gene β-actin. One-way ANOVA, Tukey post-hoc test. n = 8 per group (RT-qPCR); n = 10-12 per group (flow cytometry). *p<0.05; **p<0.01; ***p<0.001; ****p<0.0001.

### N-acetyl-cysteine improves ASD-related behaviors in *Shank3b*^-/-^ mice

In our previous work on mice lacking the ASD-relevant gene *Cntnap2*, we showed that N-acetylcysteine (NAC) may act as anti-inflammatory molecule within the cerebellum and in the periphery (12). To assess whether ASD-related behaviors may be improved in the *Shank3b* model, NAC was intraperitoneally injected once a day for 28 days into *Shank3b*^+/+^, *Shank3b*^+/-^, and *Shank3b*^-/-^ mice. At the end of the treatment, behavioral tests were performed NAC- and PBS-treated mice to assess motor (open field and rotarod test), repetitive (marble burying test), and social behaviors (3-chamber social test) (Fig. 3). As previously described (17,18), PBS-treated *Shank3b*^-/-^ mice were hypoactive and spent less time moving in the open field arena, when compared to control mice (Fig. 3a). These differences increased over time, and they were particularly evident after 20 minutes of test. In addition, *Shank3b*^-/-^ mice spent less time in the central part of the arena in relationship to the *Shank3b*^+/+^, suggesting that anxiety-like behaviors may be present in these mice (Fig. 3b). Despite this, no differences across genotypes were found after the administration of NAC. Furthermore, motor coordination was measured in NAC- and PBS-treated animals using rotarod test (Fig. 3c). Reduced time on rotarod was found in both PBS-treated *Shank3b*^+/-^ and *Shank3b*^-/-^ mice in comparison with *Shank3b*^+/+^ controls, but no changes were observed after NAC administration. Next, marble burying text was performed to investigate repetitive behaviors (Fig.3d). Again, differences between groups became evident over time and they were quantified at the end of the test (20 minutes). When the PBS-treated groups were compared, the number of buried marbles decreased in *Shank3b*^+/-^ and it was the lowest in *Shank3b*^-/-^ animals in comparison with *Shank3b*^+/+^ mice. Interestingly, a partial rescue in the phenotype was observed in both *Shank3b*^+/-^ and *Shank3b*^-/-^ animals after the injection of NAC (Fig. 3d). We finally assessed social behavior in all experimental groups using the 3-chamber social test (Fig. 3e). As expected, PBS-treated *Shank3b*^-/-^ mice showed reduced sociability index compared to *Shank3b*^+/+^ control mice. Importantly, social impairments were completely rescued in *Shank3b*^-/-^ mice administered with NAC. No differences were observed between *Shank3b*^+/-^ and *Shank3b*^+/+^ mice, in both the PBS- and NAC-treated groups. Taken together, these results show that NAC treatment could completely rescue sociability in *Shank3b*^-/-^ mice.

**Figure 3.**
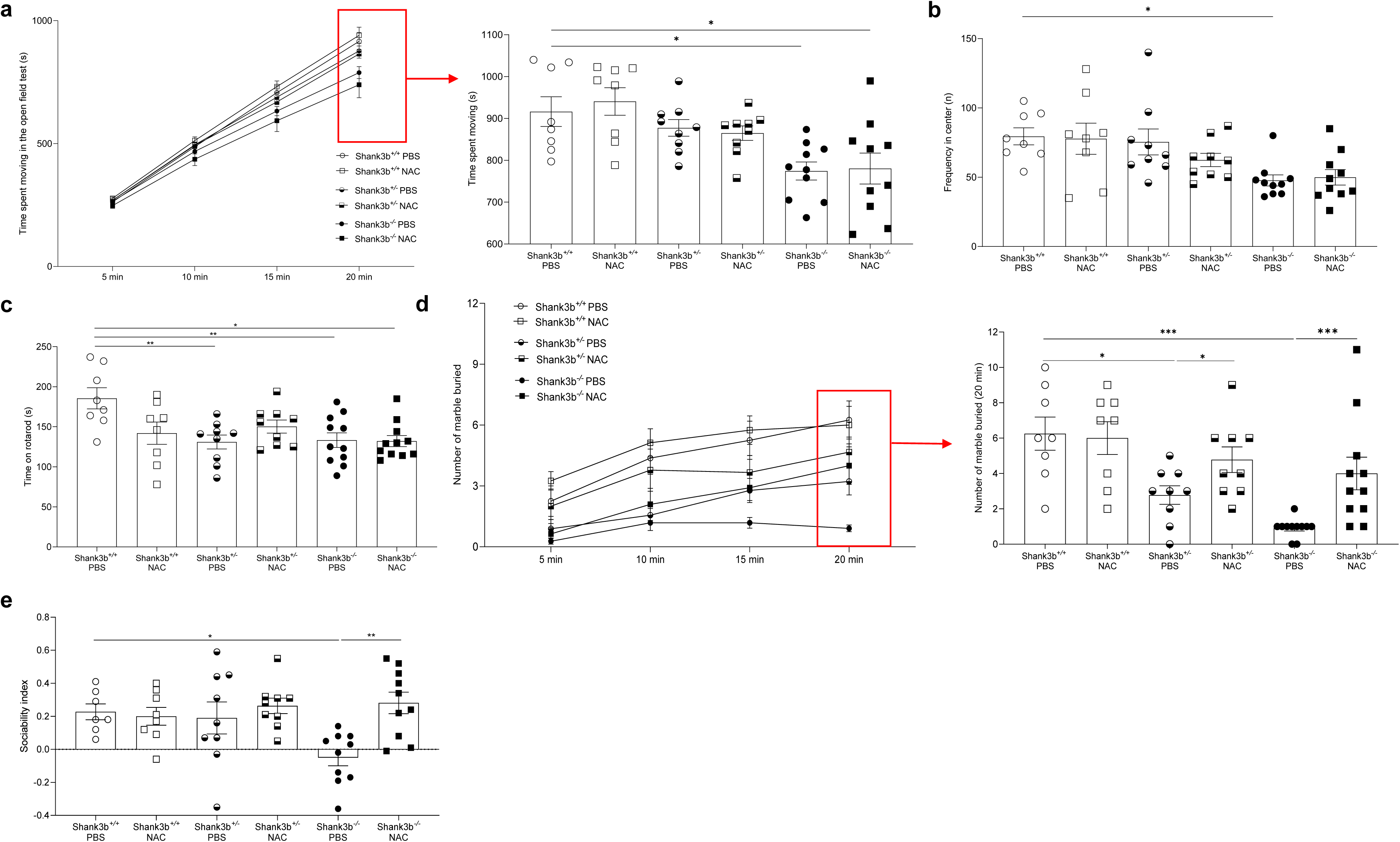
Behavioral tests in *Shank3b^+/+^*, *Shank3b^+/-^* and *Shank3b^-/-^* mice treated with NAC. Time spent moving (s) (**a**) and frequency in the center (**b**) in the open field test in NAC-treated *Shank3b^+/+^*, *Shank3b^+/-^* and *Shank3b^-/-^* mice and PBS-treated control animals. (**b**) Time on rotarod (latency to fall, s) in the rotarod test. (**c**) Number of marble buried in the marble burying test. (**d**) Sociability index (time in the sniffing zone of mouse chamber – time in the sniffing zone of empty chamber)/ total time in the sniffing zones) in the 3-chamber test. n = 8 (*Shank3b^+/+^,* PBS), n = 8 (*Shank3b^+/+^,* NAC), n = 9 (*Shank3b^+/-^,* PBS), n = 9 (*Shank3b^+/-^,* NAC), n = 9-11 (*Shank3b^-/-^,* PBS), n = 10 (*Shank3b^-/-^,* NAC). Two-way ANOVA, Tukey post-hoc test. *p<0.05; **p<0.01, ***p<0.001.

### NAC reduces pro-inflammatory impairments in the cerebellum and peripheral blood of *Shank3b*^-/-^ mice

To investigate whether improvements in ASD-related behaviors may be paralleled by decreased inflammation, pro-inflammatory molecules were quantified in the cerebellum of *Shank3b*^+/+^, *Shank3b*^+/-^, and *Shank3b*^-/-^ mice treated either with PBS or NAC (Fig.4). A decreased expression of TNF, IFNγ, IL-6, and IL-1β mRNAs was detected in both *Shank3b*^+/-^ and *Shank3b*^-/-^ mice administered with NAC, compared to controls (Fig. 4 a-d). As previously observed in *Cntnap2* mutant mice (12), the levels of these molecules (with the exception of IL-1β) was increased after NAC treatment in *Shank3b*^+/+^ mice. Similar trends were observed for CCL3, CCL5, and CCL20 chemokines, which showed a reduced mRNA expression in the cerebellum in *Shank3b*^+/-^ and *Shank3b*^-/-^ mice and increased levels in the *Shank3b*^+/+^ treated with NAC compared to their respective PBS-treated controls (Fig. 4 e-g). Furthermore, MMP8 levels were again downregulated in the NAC-treated *Shank3b*^+/-^ and *Shank3b*^-/-^ groups, while no differences were found for *Shank3b*^+/+^ mice (Fig. 4h). In addition, while p21 expression was increased in NAC-treated *Shank3b*^+/+^ and *Shank3b*^+/-^ animals, the levels of this molecule were decreased in the *Shank3b*^-/-^ group compared to their respective PBS-treated controls (Fig. 4i). In line with these observations, TNF, IFNγ, and IL-6 protein levels (respectively within CD14^+^ cells, T cells and again CD14^+^ cells) were decreased in NAC-treated *Shank3b*^+/-^ and *Shank3b*^-/-^ mice (Fig. 4 j-l). No differences were detected for *Shank3b*^+/+^ mice. These results indicate that the expression of molecules related to inflammation/damage was reduced in the cerebellum of *Shank3b*^-/-^ mice treated with NAC.

**Figure 4.**
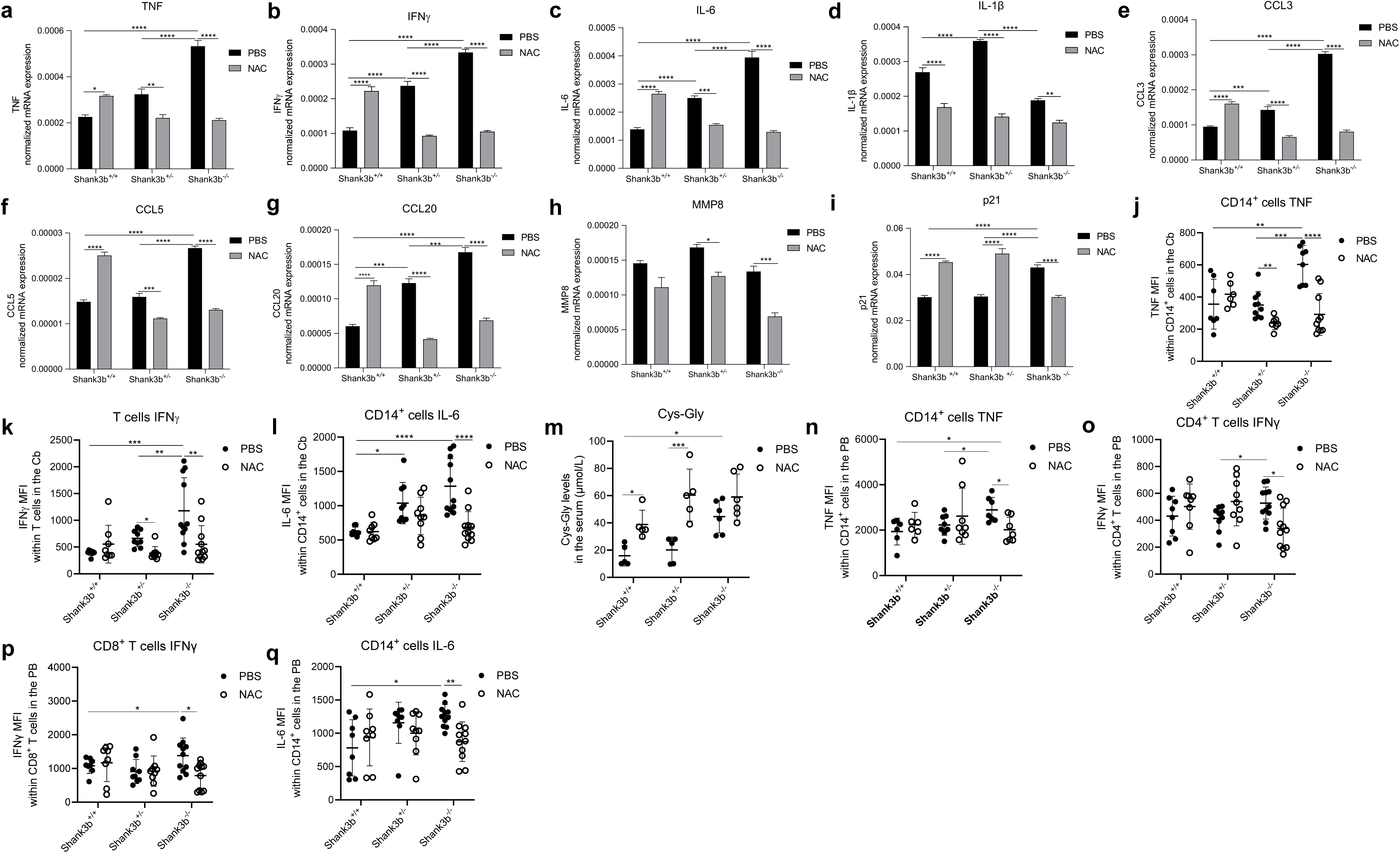
Pro-inflammatory molecules in the cerebellum of *Shank3b^+/+^*, *Shank3b^+/-^* and *Shank3b^-/-^* mice treated with NAC. mRNA expression of (**a**) TNF, (**b**) IFNγ, (**c**) IL-6, (**d**) IL-1β, (**e**) CCL3, (**f**) CCL5, (**g**) CCL20, (**h**) MMP8, and (**i**) p21 in the cerebellum of NAC-treated *Shank3b^+/+^*, *Shank3b^+/-^* and *Shank3b^-/-^* mice and PBS-treated control animals assessed using RT-qPCR. mRNA expression was normalized against the housekeeping gene β-actin. n=8 in each group. Mean fluorescence intensity (MFI) of (**j**) TNF within CD14^+^ cells, (**k**) IFNγ within T cells, and (**l**) IL-6 within CD14^+^ cells in the cerebellum measured using flow cytometry. (**m**) Cys-gly levels in the serum of PBS- and NAC-treated animals. (**n**) TNF within CD14^+^ cells, (**o**) IFNγ within CD4^+^ T cells, (**p**) IFNγ within CD8^+^ T cells, and (**q**) IL-6 within CD14^+^ cells in the peripheral blood (PB). n = 8 (*Shank3b^+/+^,* PBS), n = 8 (*Shank3b^+/+^,* NAC), n = 9 (*Shank3b^+/-^,* PBS), n = 9 (*Shank3b^+/-^,* NAC), n = 9-11 (*Shank3b^-/-^,* PBS), n = 10 (*Shank3b^-/-^,* NAC); for (m) N=5-6 in each group. Two-way ANOVA, Tukey post-hoc test. *p<0.05; **p<0.01, ***p<0.001, ****p<0.0001.

We next assessed whether NAC treatments may additionally target oxidative stress and inflammation in the PB. Plasma content of cysteinyl-glycine (Cys-Gly) in *Shank3b*^+/+^, *Shank3b*^+/-^, and *Shank3b*^-/-^ mice increased after the administration of NAC, suggesting that glutathione synthesis may be induced in these animals (Fig. 4m). When TNF expression was measured in CD14^+^ cells within peripheral blood mononuclear cells (PBMCs), decreased levels of this cytokine were found within monocytes (CD14^+^ cells) and CD4^+^ T cells (but not in CD8^+^ T cells) in NAC-treated *Shank3b*^-/-^ mice compared with PBS-treated *Shank3b*^-/-^ animals (Fig. 4n and Suppl. Fig. 3 a,b). Similarly, IFNγ expression within CD4^+^ and CD8^+^ T cells, as well as IL-6 expression within CD14^+^ cells and CD4^+^ and CD8^+^ T cells, was reduced in NAC-treated *Shank3b*^-/-^ mice (Fig. 4 n-q and Suppl. Fig. 3 c,d). No differences were observed for *Shank3b*^+/+^ and *Shank3b*^+/-^ animals. Taken together, NAC treatment rescued pro-inflammatory dysfunction in both cerebellum and PB of *Shank3b*^-/-^ animals.

### NAC counteracts immune dysfunction in the bone marrow of *Shank3b*^-/-^ **mice**

As imbalanced production of pro-inflammatory cytokines was additionally observed in the BM of *Shank3b*^-/-^ mice, the expression of molecules related to inflammation was assessed within the BM of PBS- and NAC-treated mutant and *Shank3b*^+/+^ mice (Fig. 5). The levels of all TNF, IFNγ, IL-6, and IL-1β in *Shank3b*^-/-^ were reduced after NAC administration (Fig. 5 a-d). The situation was completely different for the *Shank3b*^+/-^ group, in which NAC treatment increased IFNγ mRNA levels but no differences in the expression of the other molecules in comparison with their respective controls. Furthermore, increased IL-6 mRNA levels and no differences in the other molecules were found in *Shank3b*^+/+^ animals. Similar results were observed when mRNA expression of CCL3, CCL5, and CCL20 chemokines was assessed (Fig. 5 e-g). Following NAC treatment, CCL3 and CCL20 mRNA levels decreased in *Shank3b*^-/-^ mice, CCL20 increased in *Shank3b*^+/-^ mice, and CCL5 increased in *Shank3b*^+/+^ animals compared to their respective PBS-treated controls. Furthermore, NAC reduced MMP8 expression in the *Shank3b*^+/+^ and *Shank3b*^-/-^ groups, while increased levels were found in NAC-treated *Shank3b*^+/-^ mice. In parallel, similarly to what observed in the cerebellum, p21 mRNA expression was decreased in NAC-treated *Shank3b*^-/-^ mice, while no differences were found in the other groups. In addition, the levels of memory T cell-survival factor IL-15 did not vary between the groups (Fig. 5j). Similar results were obtained when the expression of pro-inflammatory cytokines was measured at the protein level (Fig. 5 k-n). Indeed, reduced expression of TNF within CD14^+^ cells, IFNγ in CD4^+^ and CD8^+^ T cells, and IL-6 within CD14^+^ cells was observed in NAC-treated *Shank3b*^-/-^ mice compared with the PBS-treated mice. Increased IFNγ levels in CD4^+^ and CD8^+^ T cells were present in NAC-treated *Shank3b*^+/-^ mice, while no differences were detected in the *Shank3b*^+/+^ group. Taken together, these results indicate that NAC counteracts immune dysfunction in the BM of *Shank3b*^-/-^ mice.

**Figure 5.**
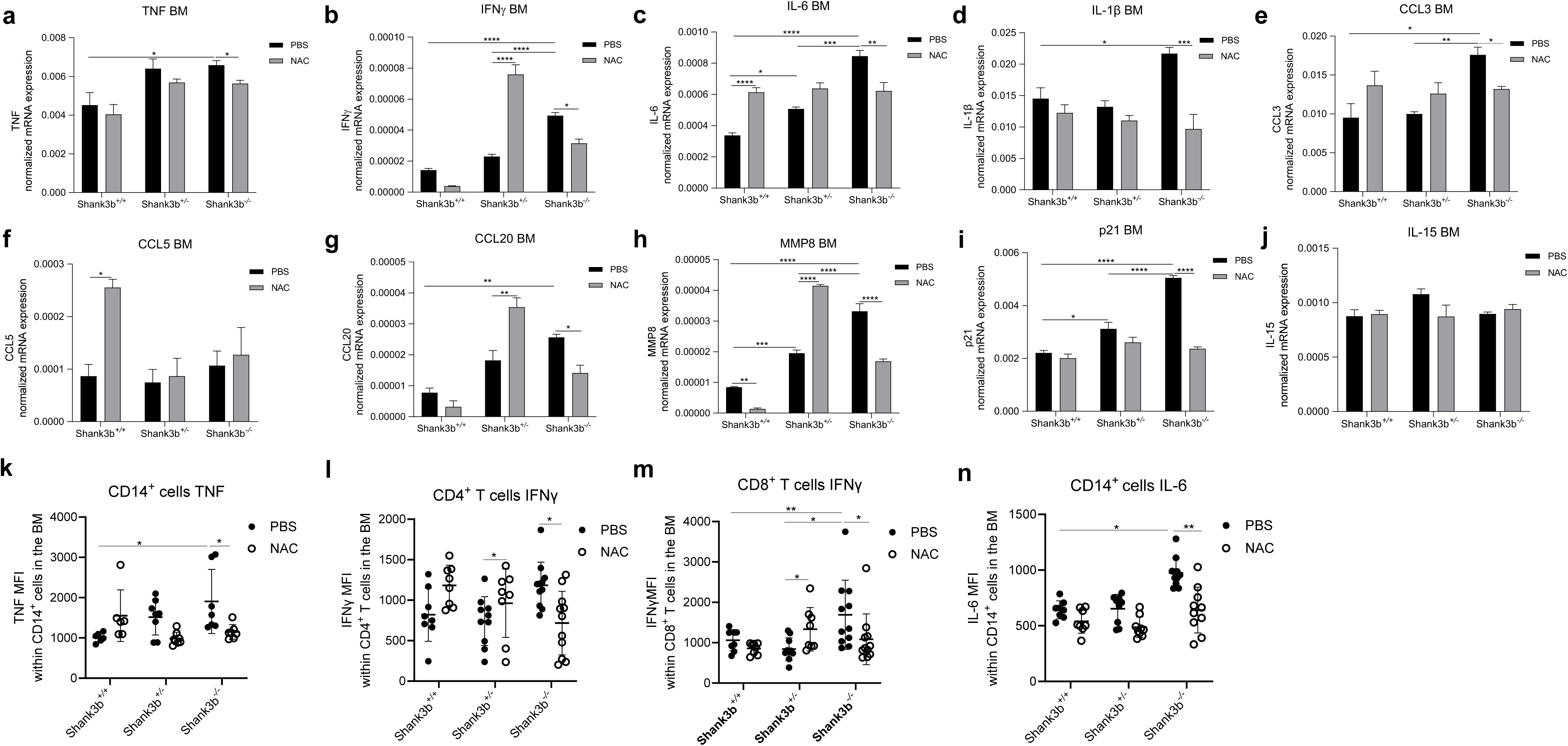
Pro-inflammatory molecules in the bone marrow of *Shank3b^+/+^*, *Shank3b^+/-^* and *Shank3b^-/-^* mice treated with NAC. mRNA expression of (**a**) TNF, (**b**) IFNγ, (**c**) IL-6, (**d**) IL-1β, (**e**) CCL3, (**f**) CCL5, (**g**) CCL20, (**h**) MMP8, (**i**) p21, and (**j**) IL-15 in the bone marrow (BM) of NAC-treated *Shank3b^+/+^*, *Shank3b^+/-^* and *Shank3b^-/-^* mice and PBS-treated control animals assessed using RT-qPCR. mRNA expression was normalized against the housekeeping gene β-actin. n=8 in each group. Mean fluorescence intensity (MFI) of (**k**) TNF within CD14^+^ cells, (**l**) IFNγ within CD4^+^ T cells, (**m**) IFNγ within CD8^+^ T cells, and (**n**) IL-6 within CD14^+^ cells in the BM measured using flow cytometry. n = 8 (*Shank3b^+/+^,* PBS), n = 8 (*Shank3b^+/+^,* NAC), n = 9 (*Shank3b^+/-^,* PBS), n = 9 (*Shank3b^+/-^,* NAC), n = 9-11 (*Shank3b^-/-^,* PBS), n = 10 (*Shank3b^-/-^,* NAC); for (m) N=5-6 in each group. Two-way ANOVA, Tukey post-hoc test. *p<0.05; **p<0.01, ***p<0.001, ****p<0.0001.

### NAC boosts pro-inflammatory processes in the spleen of *Shank3b*^-/-^ mice

Reduced pro-inflammatory responses were found in the spleen of *Shank3b*^-/-^ mice in basal conditions (Fig. 2). We therefore investigated whether NAC treatments may affect the production of pro-inflammatory molecules in spleen cells from *Shank3b*^+/+^, *Shank3b*^+/-^, and *Shank3b*^-/-^ mice (Fig. 6). Interestingly, mRNA expression of TNF, IFNγ, and IL-6 in *Shank3b*^-/-^ animals strongly increased after the NAC administration (Fig. 6 a-c). No differences were observed in *Shank3b*^+/+^ and *Shank3b*^+/-^ mice. In parallel, IL-1β levels (which were reduced in PBS-treated *Shank3b*^+/-^ mice compared with PBS-treated *Shank3b*^+/+^ controls), increased in the *Shank3b*^+/-^ group after NAC treatment (Fig. 6d). No differences were found in *Shank3b*^-/-^ animals. Furthermore CCL3, CCL5, CCL20, and MMP8 mRNAs were overexpressed in NAC-treated *Shank3b*^-/-^ mice compared with their respective PBS-treated controls (Fig. 6 e-g). Increased CCL20 mRNA expression was additionally documented for *Shank3b*^+/+^ mice, while no differences could be detected for *Shank3b*^+/-^ animals. Furthermore, p21 mRNA expression in the spleen was not affected by NAC (Fig. 6 j). IL-15 mRNA expression increased after NAC treatment in *Shank3b*^+/-^ mice (Fig. 6j). No changes were observed in the other groups. We then assessed the expression of pro-inflammatory molecules TNF, IFNγ, and IL-6 using flow cytometry (Fig. 6 k-n). Similar to the previous results, the levels of TNF within CD14^+^ cells, IFNγ in CD4^+^ and CD8^+^ T cells, and IL-6 in CD14^+^ cells increased in NAC-treated *Shank3b*^-/-^ mice (Fig. 6 l). In summary, NAC administration increases the production of pro-inflammatory molecules in the spleen of *Shank3b*^-/-^ animals, suggesting that NAC may play an important role in supporting immune responses in mutant mice.

**Figure 6.**
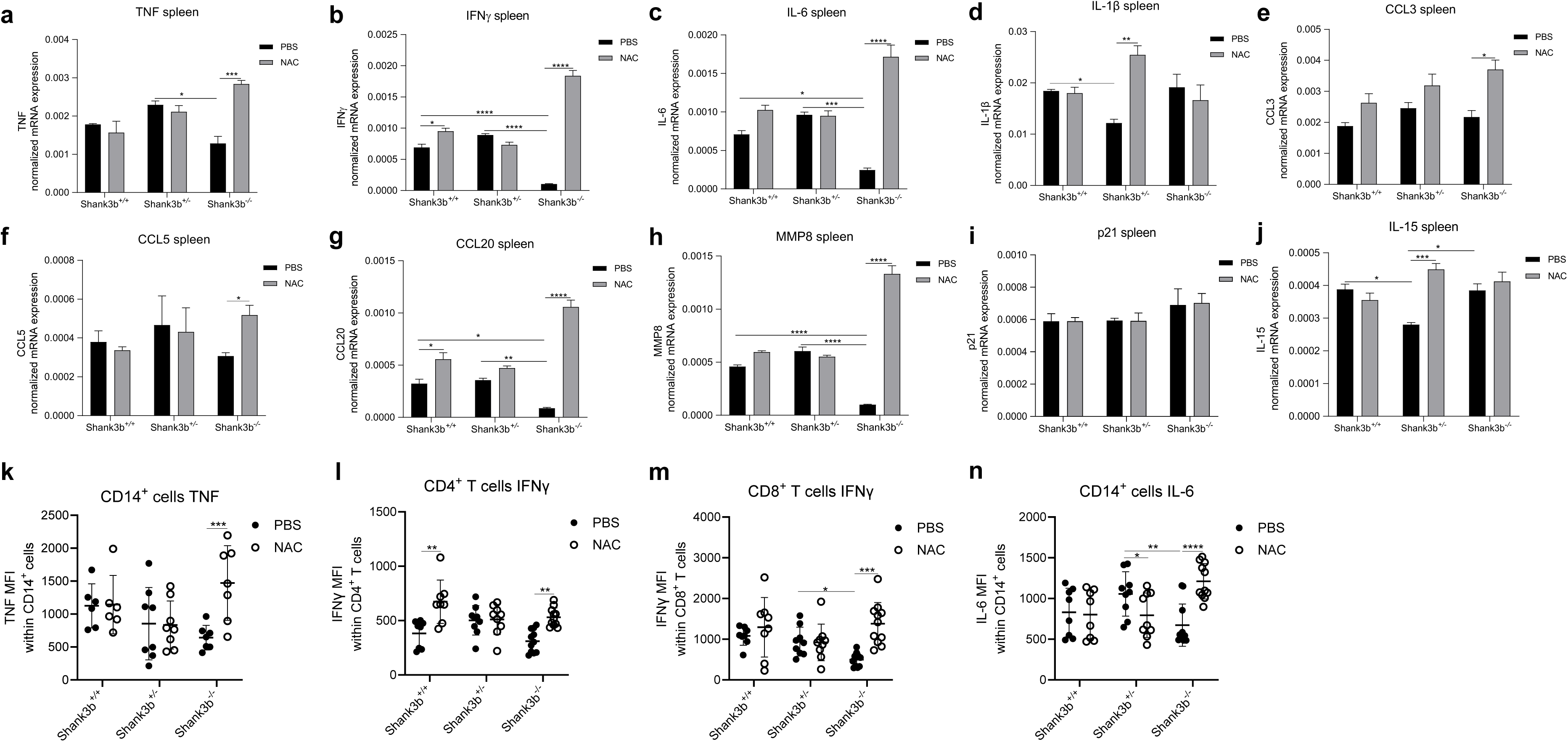
Pro-inflammatory molecules in the spleen of *Shank3b^+/+^*, *Shank3b^+/-^* and *Shank3b^-/-^*mice treated with NAC. mRNA expression of (**a**) TNF, (**b**) IFNγ, (**c**) IL-6, (**d**) IL-1β, (**e**) CCL3, (**f**) CCL5, (**g**) CCL20, (**h**) MMP8, (**i**) p21, and (**j**) IL-15 in the spleen of NAC-treated *Shank3b^+/+^*, *Shank3b^+/-^* and *Shank3b^-/-^* mice and PBS-treated control animals assessed using RT-qPCR. mRNA expression was normalized against the housekeeping gene β-actin. n=8 in each group. Mean fluorescence intensity (MFI) of (**k**) TNF within CD14^+^ cells, (**l**) IFNγ within CD4^+^ T cells, (**m**) IFNγ within CD8^+^ T cells, and (**n**) IL-6 within CD14^+^ cells in the spleen measured using flow cytometry. N = 8 (*Shank3b^+/+^,* PBS), n = 8 (*Shank3b^+/+^,* NAC), n = 9 (*Shank3b^+/-^,* PBS), n = 9 (*Shank3b^+/-^,* NAC), n = 9-11 (*Shank3b^-/-^,* PBS), n = 10 (*Shank3b^-/-^,* NAC). Two-way ANOVA, Tukey post-hoc test. *p<0.05; **p<0.01, ***p<0.001, ****p<0.0001.

## Discussion

Several recent studies have described increased levels of pro-inflammatory molecules and oxidative stress in children with ASD and mouse models (7–16). As recently proposed, inflammation may impair the maturation of vulnerable neurons, therefore supporting the onset of ASD (25). Despite this, the origin of these pro-inflammatory conditions is still unknown.

In this work, we investigated the expression of molecules related to inflammation in the cerebral cortex, hippocampus and cerebellum of the *Shank3b* mouse model of ASD. Disruption of *Shank3* has been associated with core neurodevelopmental and neurobehavioral deficits in the 22q13 deletion syndrome (Phelan-McDermid syndrome), a syndromic form of ASD (19). Previous work from our and other laboratories showed that mice lacking the *Shank3b* variant (*Shank3b^−^*^/*−*^ mice) showed pronounced ASD-related behaviors, including social impairment, self-injurious repetitive grooming, and sensory differences (17,18). Despite this, whether pro-inflammatory conditions may be present in these mice was never assessed before. As in clinical conditions *Shank3* mutations are generally present in heterozygosis, our analysis was further extended to *Shank3b^+^*^/*−*^ animals. In line with our recent observations in *Cntnap2^-/-^* mice (another mouse model of syndromic ASD; 26,12), the overexpression of pro-inflammatory molecules was specifically restricted within the cerebellum of *Shank3b^-/-^* mice (Fig.1). Indeed, *Shank3b* is highly expressed in the cerebellum (27). Overall, an intermediate level of inflammation was present in *Shank3b^+/-^* animals, suggesting that *Shank3b* gene dosage influences pro-inflammatory responses within the cerebellum. In particular, all key pro-inflammatory cytokines (TNF, IFNγ, IL-6, IL-1β) and molecules involved in immune cell migration (CCL3, CCL5 and CCL20) were upregulated in the *Shank3b^-/-^* cerebellum. While TNF was increased in CD14^+^ cells, innate immune cells which include macrophages and microglia cells (28,29), IFNγ was overexpressed in T cells. CCL3, CCL5 and CCL20 are known to be induced in the presence of pro-inflammatory molecules as they regulate the migration of immune cells to sites of inflammation (30). Altogether, these three chemokines play a determinant role in neuroinflammation as CCL3 regulate the functions of macrophages and astrocytes, (31) while CCL5 and CCL20 mainly control T cell migration (32,33). Within the cerebellum, chemokines support the pro-inflammatory profile of microglia (34). In addition, to promote the migration of immune cells, chemokines induce the expression of MMPs including MMP8, a molecule involved in tissue remodelling (35). Indeed, MMP8 within the cerebellum was highly expressed in *Shank3b^-/-^* mice. Furthermore, the cell cycle inhibitor p21, typically induced in the presence of DNA damage and considered a marker of cellular senescence (22,36), was additionally overexpressed in *Shank3b^-/-^* animals. Importantly, the mRNA expression of most of the molecules (IFNγ, IL-6, CCL3, CCL20 and p21) in *Shank3b^+/-^*mice was intermediate between *Shank3b^+/+^* and *Shank3b^-/-^*mice. In parallel, pro-inflammatory impairments were additionally detected in the PB, in both *Shank3b^+/-^* and *Shank3b^-/-^* animals.

Unexpectedly, lower mRNA levels of pro-inflammatory molecules were detected in the BM of *Shank3b^+/-^* mice compared to control *Shank3b^+/+^* mice. The situation was even more complex for *Shank3b^-/-^* mice, which showed increased levels of TNF, IL-1β, and p21 and decreased expression for IFNγ, CCL20, and MMP8. We can therefore speculate that, in the BM, the functionality of immune system may be impaired in *Shank3b^+/-^* mice, although some changes observed at the mRNA level may be at least partially “buffered” at the protein level (*i.e.* IFNγ and IL-6). Immune dysfunction may additionally be present in *Shank3b^-/-^* mice, although in this case it may be compensated by the systemic increase of pro-inflammatory molecules. Fewer differences were found in the spleen, in which the most significant differences are represented by decreased IFNγ, CCL5, CCL20 and MMP8 mRNAs in *Shank3b^-/-^* animals. In addition, antioxidant enzyme SOD3 was shown to be decreased in mutant mice in both BM and spleen.

We therefore administered the antioxidant/anti-inflammatory molecule NAC, which in our previous study was shown to target cerebellar as well as systemic inflammation in *Cntnap2^-/-^* mice Differently from *Cntnap2^-/-^* mice, no rescuing effects were observed on motor deficits in *Shank3b^-/-^* animals as detected by the open field and rotarod tests. This apparent inconsistency may be explained by a key difference between the *Cntnap2* and *Shank3b* models, as *Cntnap2^-/-^* mice are *hyperactive* while *Shank3b^-/-^* mice are *hypoactive*. Thus, NAC may be more effective in counteracting hyperactivity rather than hypoactivity, as previously observed (37). In line with previous studies (38), *Shank3b^+/-^*and *Shank3b^-/-^* mice buried a lower number of marbles in the marble burying test compared to control mice. Interestingly, NAC restored this phenotype, indicating that it may at least partially rescue hypoactivity in *Shank3b^-/-^* mice. Most importantly, NAC administration was able to rescue sociability deficits in *Shank3b^-/-^*animals. These results were accompanied by decreased levels of pro-inflammatory molecules in the cerebellum, as all molecules related to inflammation were reduced in both *Shank3b^+/-^* and *Shank3b^-/-^*mice. Similarly to our previous observations in the *Cntnap2* model (12), an overall increase in the expression of pro-inflammatory molecules was observed in *Shank3b^+/+^* mice treated with NAC; this might be caused by “antioxidative stress”, a stressful condition supported by the excessive elimination of oxygen radicals (39,40). It is however worth noting that despite this pro-inflammatory activity of NAC, we did not observe social impairment in the NAC-treated *Shank3b^+/+^* group.

Similar results were detected in the PB. In this context, increased cys-gly levels (indicating glutathione synthesis) were found in the serum of all NAC-treated groups. Thus, in all animals, NAC may effectively be converted into glutathione, although the effects of the drug may be achieved also in a glutathione-independent manner (41). In the PB, all pro-inflammatory molecules of interest were decreased in *Shank3b^-/-^*mice, while only some of them were reduced in *Shank3b^+/-^* animals. Intriguingly, treatments by themselves increased levels of inflammation in the BM of *Shank3b^+/-^* and *Shank3b^-/-^* mice, as in animals treated with vehicle the expression of all key pro-inflammatory cytokines were increased when compared with *Shank3b^+/-^* and *Shank3b^-/-^*mice in basal conditions (Figures 1 and 2). This might be caused by a reduced capability of mutant mice to counteract stress. Overall, NAC administration partially boosted immune functions in the BM of *Shank3b^+/-^* mice, as IFNγ was increased in *Shank3b^+/-^* animals while some molecules (IL-6 and CCL5) were additionally induced in the *Shank3b^+/+^* group. In parallel, the levels of most pro-inflammatory molecules were reduced in NAC-treated *Shank3b^-/-^*when compared to their PBS-treated counterpart. This was accompanied by decreased levels of p21. Thus, in *Shank3b^-/-^* animals, the administration of NAC may “balance” the levels of pro-inflammatory molecules and counteract damage.

The spleen represents the largest lymphoid organ, and the production of pro-inflammatory molecules by immune cells within the spleen exerts a vital role in immunity. Thus, reduced levels of cytokines in this organ are generally associated with immune dysfunction (42). Interestingly, our results suggest that immune dysfunction observed in *Shank3b^-/-^* animals could be counteracted by NAC, as the production of all key pro-inflammatory molecules was strongly increased. Some changes were additionally documented for *Shank3b^+/+^*and *Shank3b^+/-^* mice.

In summary, this study contributes to strengthen the hypothesis that pro-inflammatory dysfunction within the cerebellum may be associated with ASD-related behaviors. In addition, we showed that immune dysfunction extends to the PB, BM, and spleen, and it can be counteracted after the administration of NAC. Although these differences were clearer in *Shank3b^-/-^* mice, a certain degree of immune dysfunction was additionally present in *Shank3b^+/-^* animals, and NAC was effective also in this group. This represents an important observation which may be considered for developing novel strategies of intervention for the clinical setting. Furthermore, this study underlines the importance to address ASD as a “systemic disorder”, and not only as a condition affecting the brain only.

## Materials and Methods

### Animals

All experimental procedures were approved by the Animal Welfare Committee of the University of Trento and Italian Ministry of Health (protocol n.922/2018-PR and n. 547/2021-PR), in accordance with the European Community Directive 2010/63/EU. Mice were housed following a 12 h light/dark cycle with food and water available *ad libitum*, taking care to minimize animal’s pain and discomfort. Male and female *Shank3b^-/-^, Shank3b^+/-^, Shank3b^+/+^* age-matched adult littermates (3–5 months old; weight 25–35 g) obtained from heterozygous mating were used. Numbers of mice used for each experiment are reported in figure legends.

### Tissue harvesting

Brain samples used for RT-PCR were dissected and frozen in dry ice. Cerebella used for flow cytometry experiments were homogenized using a homogenizer immediately after dissections, and single-cell suspension were prepared using Falcon 70μm cell strainer (Corning). Approximately two hundred μl of PB was harvested from each mouse and collected in a heparinized tube. PBMCs were isolated using a Ficoll-Hypaque density gradient (Sigma-Aldrich). BM cells were obtained after flushing femurs and tibias with PBS. Spleen samples were directly smashed through Falcon 70 µm cell strainers. After the isolation, cerebellum, PBMCs, BM and spleen cells were washed once with RPMI 1640 (Sigma-Aldrich) and resuspended in complete medium (RPMI 1640 supplemented with 10% fetal calf serum, FCS, 100 U/mL penicillin and 100 μg/mL streptomycin; Sigma-Aldrich and Invitrogen respectively).

### RNA isolation and quantitative RT-PCR (qRT-PCR)

Total RNAs were extracted from cerebral cortex, hippocampus, cerebellum, BM, and spleen samples of adult *Shank3b^-/-^, Shank3b^+/-^, Shank3b^+/+^* mice using RNeasy Plus Mini Kit (Qiagen), and retro-transcribed to cDNA as reported in our previous work (12,43). qRT-PCR was performed in a Bio-Rad C1000 Thermal Cycler, using the PowerUp™ SYBR™ Green Master Mix (Applied Biosystems). Primers (Eurofins Genomics) were designed on different exons to avoid amplification of genomic DNA. Sequences of primers using for the study are shown in TableS1. CFX3 Manager 3.0 (Bio-Rad) software was used to perform expression analyses. Mean cycle threshold (Ct) values from triplicate experiments were calculated for each gene of interest and the housekeeping gene β actin, and afterwards corrected for PCR efficiency and inter-run calibration. The expression of each mRNA of interest (normalized against β actin) was compared from triplicate experiments performed on RNA pools from 8-10 samples for each group.

### Flow cytometry

Flow cytometry was used to quantify cytokine levels and immune cell populations in the cerebellum, PB, BM and spleen. Immunofluorescence surface staining was performed by adding a panel of directly conjugated antibody to freshly prepared cells. To assess the expression of cytokines, cells were incubated with 30 ng/mL phorbol 12-myristate 13-acetate (PMA) and 500 ng/mL ionomycin in the presence of 10 mg/mL brefeldin A (BFA) (all molecules from Sigma-Aldrich) for 4 h at 37 °C. After surface staining, cells were permeabilized using the Cytofix/Cytoperm kit (BD Biosciences) and incubated with intracellular antibodies. Labeled cells were measured using a LSR Fortessa (BD Biosciences) flow cytometer available at the Institute for Biomedical Aging Research, University of Innsbruck. Data were analyzed using Flowjo software. The antibodies used in the experiments are shown in TableS2.

### NAC treatment and behavioural tests

*Shank3b^-/-^, Shank3b^+/-^, Shank3b^+/+^*mice were injected intraperitoneally with 50mg/kg NAC (Sigma-Aldrich) resuspended in PBS or vehicle (PBS) for 28 consecutive days as previously done (12).

#### Behavioral tests

Open field, rotarod, Marble burying and 3-chamber social tests were performed during the last 5 days of NAC treatment of *Shank3b^-/-^, Shank3b^+/-^, Shank3b^+/+^* mice. In all behavioral tests, male and female mice were habituated and tested separately to avoid experimental noise.

#### Open field test

Open field test was performed to measure motor activity of mice. Animals were placed in empty open field arena (40 cm × 40 cm × 40 cm) and allowed to freely explore it for 20 min. Sessions were recorded by a camera located over the arena and mice were automatically video tracked using the software EthoVisionXT (Noldus). Time spent moving and frequency in the center of the arena were analysed.

#### Rotarod test

Cerebellar-associated motor coordination was assessed using rotarod test (44). A habituation phase conducted at constant speed of 4 rpm was performed in the two days before the test. Experimental phase consisted in two trials in which the rotation speed was increased from 4 rpm to 64 rpm. Falling mice landed on a metallic platform that was connected to a timer showing the time spent on the rotating rod. The average time spent on the rod (latency to fall) during the second trial was measured in all experimental groups, and it was used as quantitative indicator of the motor ability of the mice to stay balanced on the rotating accelerating rod.

#### Marble burying test

Marble burying test was performed as described previously (38). Tested mouse was placed in a cage containing bedding at a depth of 2 cm. 20 black glass marbles were arranged on top of the bedding. The mouse was placed in the center of the cage for a 20-min exploration period, under 15 lux illumination and pictures were taken every 5 minutes. Number of marbles at least 50% covered by bedding were scored as buried.

#### Three-Chamber social test

The three-Chamber social test was used to assess the social behaviors of mice (45). The apparatus consisted of a plexiglass rectangular box (60×40×22(h) cm, each chamber 20×40×22(h) cm) with grey coloured walls and with removable panels separating the box into three chambers. On the four days before the beginning of the experimental phase, mice underwent a habituation phase in which they were placed in the three-chamber apparatus and allowed to freely explore it for 10 min. The experimental phase consisted of a 5-minutes habituation session, a 10-minutes exploration session and the sociability test, in which one unfamiliar mouse was placed into a wire cylindrical cage (20 cm in height, 10 cm bottom-diameter, 1cm bars spaced) in one of the up corner of an external chamber. An identical empty wire cage was placed in the opposite external chamber. The tested mouse was allowed to freely interact with the unfamiliar mouse and with wire cages for 10 minutes. The sociability index was calculated as the ration between the time spent in the social chamber and the total time spent in the external chambers. Trials were recorded by an overhead camera placed over the three-chamber apparatus. Mice were automatically video tracked using EthoVisionXT.

### Thiols determination

Serum Cysteinyl-Glycine (Cys-Gly) levels were analyzed as previously reported (46).

### Statistics

Statistical analyses were performed with GraphPad Prism 8.0 software, using one-way ANOVA and two-way ANOVA followed by Tukey post-hoc test, with the level of significance set at p<0.05.

## Author contribution

LP designed and supervised the study, provided funding, performed molecular and behavioral experiments, analysed data, and wrote the manuscript; EC performed molecular and behavioral experiments and analysed data; LB performed behavioral experiments and contributed to NAC treatments, GMD performed molecular experiments; GC contributed to NAC treatments; AP performed the cys-gly determination in the plasma and analysed data; BW contributed to the study design and edited the manuscript; YB supervised and designed the study, provided funding, and edited the manuscript.

## Acknowledgements

The authors thank the technical and administrative staff of CIMeC, University of Trento, for excellent support. CIBIO Department Core Facilities (University of Trento) and the Biooptical Centre Facility of the Institute for Biomedical Aging Research (University of Innsbruck) provided the infrastructures and assistance to perform flow cytometry experiments.

---

**Figure S1.**
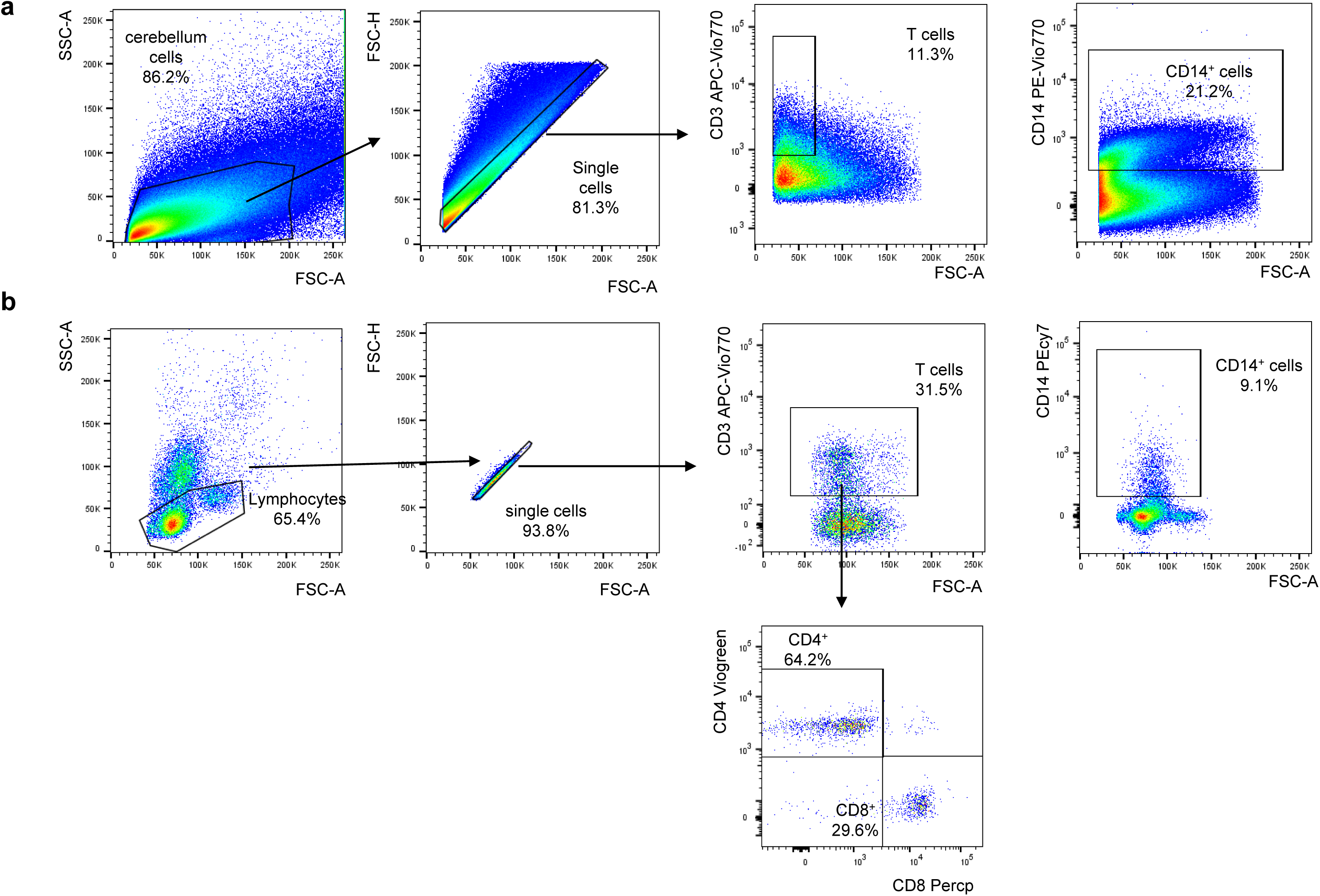
Gating strategy used for the flow cytometry experiments. **(a)** Gating strategy used to define (**a**) T cells and CD14^+^ cells within cerebellar cells, and (**b**) T cells (CD8^+^ and CD4^+^), and monocytes (CD14^+^ cells) within PBMCs.

**Figure S2.**
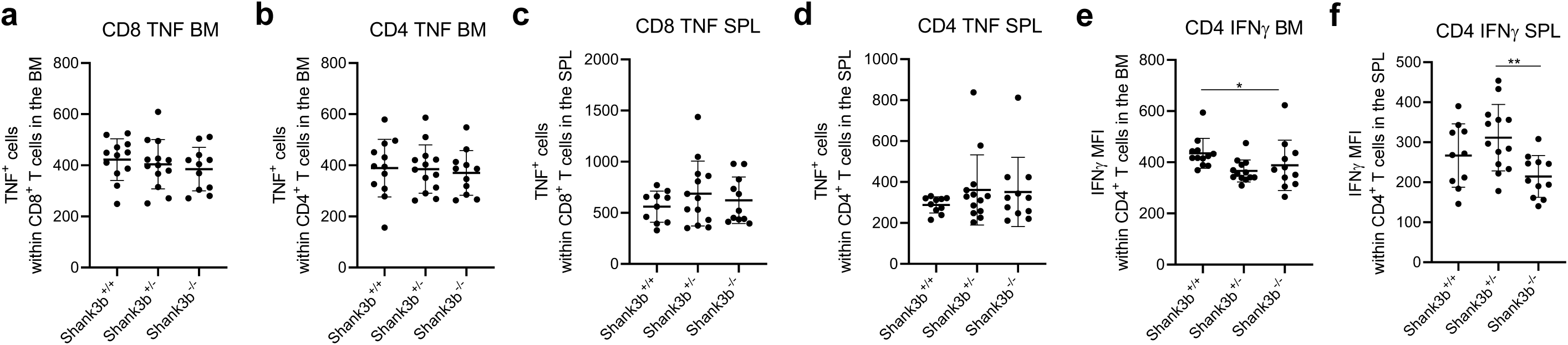
Pro-inflammatory molecules within T cells in the bone marrow and spleen of *Shank3b^+/+^*, *Shank3b^+/-^* and *Shank3b^-/-^* mice. Mean fluorescence intensity (MFI) of TNF within (**a**) CD8^+^ T cells, and (**b**) CD4^+^ T cells in the BM; MFI of TNF within (**c**) CD8^+^ T cells, and (**d**) CD4^+^ T cells in the spleen; MFI of IFNγ within CD4^+^ T cells in the (**e**) BM and (**f**) spleen. One-way ANOVA, Tukey post-hoc test. ***p<0.001. n = 10-12 per group.

**Figure S3.**
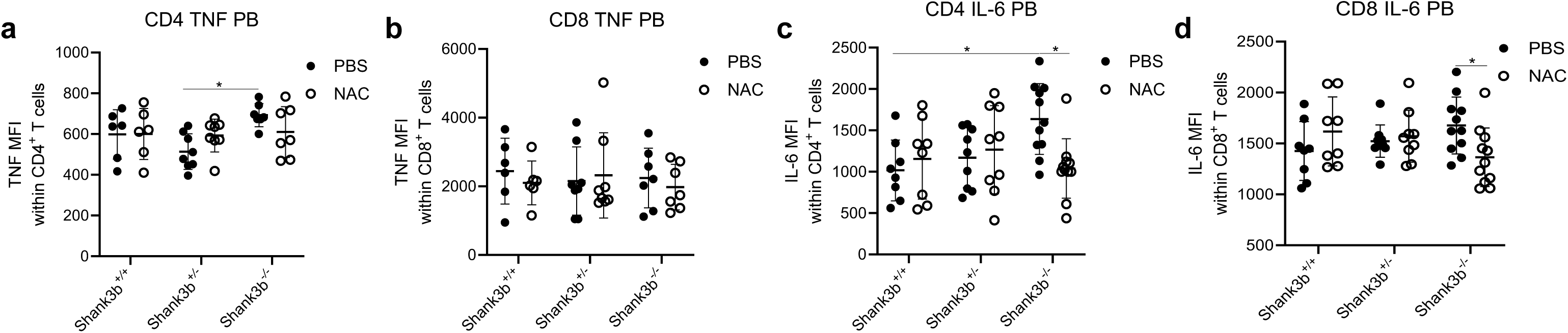
Pro-inflammatory molecules in the PB of *Shank3b^+/+^*, *Shank3b^+/-^* and *Shank3b^-/-^*mice treated with NAC. Mean fluorescence intensity (MFI) of TNF within (**a**) CD4^+^ T cells, (**b**) CD8^+^ T cells, and IL-6 MFI within (**c**) CD4^+^ T cells, and (**d**) CD8^+^ T cells in the PB of NAC-treated *Shank3b^+/+^*, *Shank3b^+/-^* and *Shank3b^-/-^* mice and PBS-treated control animals. Two-way ANOVA, Tukey post-hoc test. *p<0.05. n = 10-12 per group.

